# Stromal depletion by TALEN-edited universal hypoimmunogenic FAP-CAR T cells enables infiltration and anti-tumor cytotoxicity of tumor antigen-targeted CAR-T immunotherapy

**DOI:** 10.1101/2023.02.27.529938

**Authors:** Shipra Das, Julien Valton, Philippe Duchateau, Laurent Poirot

**Author notes:** **Correspondence:** Shipra Das.

## Abstract

Adoptive cell therapy based on chimeric antigen receptor-engineered T (CAR-T) cells has proven to be lifesaving for many cancer patients. However, its therapeutic efficacy has so far been restricted to only a few malignancies, with solid tumors proving to be especially recalcitrant to efficient therapy. Poor intra-tumor infiltration by T cells and T cell dysfunction due to a desmoplastic, immunosuppressive microenvironment are key barriers for CAR T-cell success against solid tumors.

Cancer-associated fibroblasts (CAFs) are critical components of the tumor stroma, evolving specifically within the tumor microenvironment in response to tumor cell cues. The CAF secretome is a significant contributor to the extracellular matrix and a plethora of cytokines and growth factors that induce immune suppression. Together they form a physical and chemical barrier which induces a T cell-excluding ‘cold’ TME. CAF depletion in stroma rich solid tumors can thus provide an opportunity to convert immune evasive tumors susceptible to tumor-antigen CAR T-cell cytotoxicity.

Using our TALEN-based gene editing platform we engineered non-alloreactive, immune evasive CAR T-cells (termed UCAR T-cells) targeting the unique CAF marker Fibroblast Activation Protein, alpha (FAP). In an orthotopic mouse model of triple-negative breast cancer (TNBC) composed of patient derived-CAFs and tumor cells, we demonstrate the efficacy of our engineered FAP UCAR T-cells in CAF depletion, reduction of desmoplasia and successful tumor infiltration. Furthermore, while previously resistant, pre-treatment with FAP UCAR T-cells now sensitized these tumors to Mesothelin (Meso) UCAR T-cell infiltration and anti-tumor cytotoxicity. Combination therapy of FAP UCAR, Meso UCAR T cells and the checkpoint inhibitor anti-PD-1 significantly reduced tumor burden and prolonged mice survival. Our study thus proposes a novel treatment paradigm for successful CAR-T immunotherapy against stroma-rich solid tumors.

## Introduction

Since its inception in 2012^1^, clinical application of autologous chimeric antigen receptor (CAR)^2^ T cell therapies for hematological malignancies have witnessed remarkable success and advancement^3–6^. However, CAR T-cell therapy against solid tumors, which comprise about 90% of the total cancer landscape^7^, has encountered very limited success. Notably, the clinical efficacy of CAR T-cell trials against the top three widely expressed solid tumor targets namely mesothelin, GPC3 and mucin-1 (MUC1) have so far been met with low to middling success^8^.

This can be attributed to several challenging properties inherent to most solid tumors^9–11^. Indeed, unlike most hematological malignancies, the bulk of solid tumors have a distinct heterotypic microenvironment characterized by a desmoplastic stromal compartment and a multi-cellular immune landscape. This manifests into a fibrotic, collagen-rich network which forms a physical barrier against T cell infiltration and in most instances an immunosuppressive signature that induces T cell exhaustion and dysfunction. A significant proportion of solid cancers including ovarian, breast, colorectal and pancreatic adenocarcinomas represent such cold tumors and demonstrate a primary resistance to immune checkpoint inhibition (ICI) therapies as well as adoptive cell therapies. While tumor microenvironment (TME) properties pose complications for the patient’s own immune defense mechanisms as well as adoptive cell therapies, high antigen heterogeneity and the risk of ‘on-target off-tumor’ cytotoxicity due to sparsity of tumor-specific antigens (TSA) are additional hurdles for CAR T-cell therapy success^9, 10^. Current efforts are therefore focused on innovating CAR T-cell treatment strategies that can overcome these obstacles.

A critical component of the TME that contributes to its immune-evasive properties are cancer-associated fibroblasts (CAFs), a subpopulation of quiescent fibroblasts, which evolve specifically in the tumor milieu in response to neoplasm-derived paracrine cues. In addition to collagen-rich extracellular matrix proteins, the CAF secretome includes cytokines and growth factors like TGF-β and CXCL12, that promote immunosuppression. Together, these physical and chemical barriers foster T-cell exclusion and exhaustion^14, 15^. CAF-rich tumors are thus often ‘cold’ nonimmunogenic tumors with a T-cell deficient immune desert signature landscape^16, 17^ Corroboratively, over 90% of epithelial cancers including breast, colorectal, pancreatic and lung adenocarcinomas express the CAF-specific surface marker, fibroblast activation protein-α (FAP)^18^, that correlates with both poor prognosis and response to immunotherapies^19, 20^. Fortunately, other than a small fraction of cells in the proliferative stage of placenta and uterine stroma, embryonic tissue and multipotent bone marrow stromal cells, FAP is not expressed in healthy adult tissues^21^. Therefore, several FAP-targeting approaches are under development to target these pro-tumorigenic CAFs^22–25^. A significant revelation of these studies is that CAF ablation increases intra-tumoral T cell-infiltration in addition to reprogramming the TME to an immune reactive milieu^26, 27^ These observations unveil a whole avenue of targeting ‘cold’ tumors using adoptive cell therapies by first priming the TME with CAF-targeting approaches.

Here, we evaluated the effect of FAP CAR T-cell pre-treatment on therapeutic efficacy of CAR T-cells targeting tumor-antigens currently under investigation in clinical trials against solid tumors^8, 9^. For this purpose, we selected the tumor associated antigen (TAA) Mesothelin which is overexpressed in most solid tumors including mesothelioma and large sub-sets of ovarian, breast, pancreatic and lung adenocarcinomas.^28^. Of these, mesothelin over-expression in triple-negative breast cancer (TNBC) was of primary interest to us^29, 30^, given (i) the CAF rich TME that correlates with poor prognosis^31, 32^, and (ii) the urgent need for development of new therapies due to current lack of targeted and effective treatment options. Furthermore, much like other TAAs, while prominently expressed in tumors, mesothelin is also detected on normal peripheral tissues^21, 33^. Subsequently, non-alloreactive CAR T-cells with their finite persistence in patients can be a relatively safer immunotherapeutic approach against these tumors.

Thus, using a non-alloreactive cell therapy approach and a physiologically relevant mouse tumor model of triple-negative breast cancer, we demonstrate the pre-clinical efficacy of FAP CAR-T pre-treatment to reprogram the ‘cold’ TME and make the tumor susceptible to subsequent Mesothelin CAR T cytotoxicity. Overall, our study proposes a novel treatment paradigm of combination CAR T-cell therapy that can be extended to most stroma-rich cold tumors using tumor-antigen targeting CAR T-cells currently under development.

## Materials and Methods

### Primary cells, cell lines and cell culture

Cryopreserved human PBMCs were acquired from ALLCELLS (# PB006F). PBMCs were cultured in CTS OpTmizer media (obtained from Gibco, # A1048501), containing IL-2 (Miltenyi Biotec, # 130-097-748), human serum AB (Seralab, # GEM-100-318), and CTS Immune Cell SR (Gibco, # A2596101). Human T Cell TransAct (Miltenyi Biotec t# 130-111-160) was used to activate T cells. PBMCs were cryopreserved in 90% albumin/10% DMSO.

HCC70-GFP were engineered from HCC70 cells (ATCC, # CRL-2315) using an in house rLV encoding NanoLuc_T2A_EGFP construct and AMSbio (# LVP323-PBS), respectively, using the manufacturer’s protocols. TNBC patient-derived CAFs were obtained from BioIVT (cancer fibroblasts #233831P1). All cell lines and CAFs were maintained in DMEM supplemented with 10% heat-inactivated FBS in 5% CO2 at 37°C.

### UCAR T-cell generation and expansion

Briefly, PBMCs were thawed, washed, resuspended, and cultivated in CTS OpTmizer complete media (reconstituted CTS OpTmizer, 5% human AB serum, 20□ng/mL IL-2). One day later, the cells were activated with Human T Cell TransAct (25□μL of beads/10^6^ CD3 positive cells) and transduced with recombinant lentiviral vectors (Flash Therapeutics) at a multiplicity of infection (MOI) of 15 for FAP CAR and 10 for Mesothelin CAR T-cells, in retronectin coated culture vessels (Takara Bio USA Inc, #T100B). Cells were transduced at a concentration of 2×10^6^ cells/ml in full media with Transact and cultured at 37°C in the presence of 5% CO2 for 3 days. The cells were then split into fresh complete media and transfected the next day according to the following procedure. On the day of transfection, the cells were washed twice in Cytoporation buffer T (BTX Harvard Apparatus, Holliston, Massachusetts), and resuspended at a final concentration of 28□×□10^6^ cells/mL in the same solution. The cellular suspension (5□×□10^6^ cells) was mixed with 5□μg mRNA encoding each TRAC TALEN arm and 5 μg of mRNA encoding each arm of B2M TALEN in a final volume of 2000□μl. The cellular suspension was transfected in 0.4□cm gap cuvettes using Pulse Agile technology. The electroporation consisted of two 0.1□mS pulses at 2000□V/cm followed by four 0.2□mS pulses at 325□V/cm. Electroporated cells were transferred to a 12-well plate containing 2□mL. of prewarmed OpTmizer media (supplemented with human serum and IL-2) and incubated at 37°C for 15□min and then transferred to 30°C overnight. Next day, cells were seeded at a density of 10^6^ cells/mL in complete OpTmizer media and cultivated at 37□°C in the presence of 5%□CO2. On day 8 post thawing, the cells were resuspended in fresh complete medium supplemented with 20□ng/mL IL-2 and 5% CTS Immune Cell SR. The cells were seeded in GREX10 (Wilson Wolf, #80040S) at 0.125□×□10^6^ cell/ml and cultivated in the same media according to the manufacturer’s guidelines.

### *In vitro* T-cell antitumor activity assays

To assess cytotoxicity of FAP UCAR T-cells against CAFs, CAFs were fluorescently labelled with CFSE (0.5 mM) according to the manufacturer’s instructions (Invitrogen, #C34554) and co-incubated at 37°C, 5% CO2 with either mock or FAP UCAR T-cells at different CAF: CAR^+^ T cell ratios. After one day of co-incubation, CFSE-labelled CAFs were harvested and stained with fixable viability dye (eBioscience, #65-0865-14) and analyzed by flow cytometry on a FACSCanto II cytometer (BD). CFSE-CAFs positive for the viability dye were quantitated as dead cells and percentage of CAF lysis was determined for every condition.

To assess UCAR T-cell cytotoxicity against Tumor-CAF spheroids, on Day 0, 10^4^ triple-negative breast tumor cells HCC70, transduced to express GFP (HCC70-GFP) were seeded either alone or with TNBC-derived CAFs at a 1:1 ratio on low adherence 96-well round bottom plates (Thermo Fisher # 174925), in DMEM+10%FBS media. Tumor cells and CAFs were given 3 days to organize themselves into spheroids. On day 3, UCAR T-cells were added to the spheroids at tumor cell: CAR^+^ T cell ratio of 1:5. 72 h post co-incubation with UCAR T-cells, spheroids were imaged on Incucyte ZOOM live-cell imaging and analysis platform (Sartorius). GFP signal per spheroid was quantitated using the Incucyte Basic Analysis Software as average green object integrated intensity (GCU x um^2^). GFP intensity of control untransduced T-cell (UT) treated spheroids was normalized to 100% survival and used to calculate survival of UCAR T-cell treated spheroids.

### IFNγ secretion assay

The levels of INFγ were evaluated in supernatants obtained from 0.625:1 and 1.25:1 effector:target ratio overnight co-cultures of CAR T-cells:CAFs using the Human IFN-Gamma Quantikine Kit (R&D Systems) following manufacturer’s instructions. As positive control, CAR T-cells were activated with Ionomycin and PMA).

### Flow cytometry

For *in vitro* cell cultures, cells in U-bottom 96-well plate were spun down and washed with PBS (150 □μL/well) at 300□×□*g* for 2□min. Prior to surface staining, cells were stained with Fixable Viability Dye eFluor 450 or eFluor 780 (eBiosciences) according to the manufacturer’s instructions. The cells were then stained with antibodies diluted in FACS buffer (2% FBS□+□5□mM EDTA□+□0.05% azide in PBS, 50□μL/well) for at least 30□min in the dark at 4□°C. Cells were washed with PBS (150□μL/well), spun at 300×*g* for 2□min, and resuspended in fix buffer (4% paraformaldehyde in PBS, 100□μL/well). Sample collection was performed on a FACSCanto II cytometer (BD) or NovoCyte Penteon flow cytometer (Agilent), and data were analyzed using FlowJo V.10.6.1 (Treestar) or NovoExpress V.1.5.6 respectively.

For tumor samples, tumor tissue was chopped finely with a razor in 5 ml Accutase (Biolegend, #4232201) and incubated at 37°C water bath for 30 minutes. Digested tumor suspension was passed through 100 um strainer (Corning) and filtrate was spun at 300 x g for 10 min. Cell pellet was subsequently stained for flow cytometry as described above. Mouse spleens were processed by crushing the spleen in 5 ml PBS+2%FBS. Cell suspension was spun at 300 x g for 10 min. Cell pellet was suspended in 1X RBC lysis buffer (eBioscience, # 00-4300-54) for 5 min at room temperature and then filtered through a 70 um-strainer (Corning). Filtrate was spun at 300 x g for 7 min and cell pellet was subsequently stained for flow cytometry as described above. Staining was performed with the following antibodies:

| Differentiation |  |  |  |
| --- | --- | --- | --- |
| Antigen | Fluorochrome | Company | Catalog No. |
| CD45RA | APCCy7 | Miltenyi Biotec | 130-113-363 |
| CD62L | PE | Miltenyi Biotec | 130-113-625 |
| CD4 | PE-Vio770 | Miltenyi Biotec | 130-113-227 |
| CD8 | BV510 | BD Bioscience | 563919 |
| CD95 | PerCP-Cy5.5 | Biolegend | 305630 |
| Viability | e780 | Invitrogen | 65086514 |
| UCAR T Phenotype |  |  |  |
| FAP Protein | FITC | AcroBiosystems | FAP-HF263 |
| Mesothelin protein | PE | AcroBiosystems | MSN-HP2H8 |
| **TCRa/b** | APC | Miltenyi Biotec | 130-098-859 |
| **HLA-ABC** | VioBlue | Miltenyi Biotec | 130-101-459 |
| **Tumor analysis** |  |  |  |
| **Antigen** | **Fluorochrome** | **Company** |  |
| **hCD45** | PE | Miltenyi | 130-110-632 |
| **mCD45** | BV-421 | BD Biosciences | 560501 |
| **CD4** | PE-Vio770 | Miltenyi Biotec | 130-113-227 |
| **CD8** | BV510 | BD Bioscience | 563919 |
| **hFAP** | 488 | R&D Systems | MAB9727-100 |
| **hMesothelin** | APC | R&D Systems | FAB32652P |

### Mice and animal procedures

All procedures involving animals were approved by The Mispro Institutional Animal Care and Use Committee and were performed in accordance with the guidelines of the PHS (Public Health Service) Policy on Humane Care and Use of Laboratory Animals, OLAW (Office of Laboratory Animal Welfare), and the USDA (United States Department of Agriculture) AWA (Animal Welfare Act). Experimental/control animals were co-housed.

All experiments were performed on 8-weeks-old, female NOD.Cg-Prkdcscid Il2rgtm1Wjl/SzJ (NSG) mice obtained from The Jackson Laboratory (Stock # 005557). Animals were housed in SPF animal facility. Mouse room light cycles were on a 12□h on/off (on from 6□am to 6□pm and off from 6□pm to 6□am). temperature reading was maintained between 68 and 79□F, and humidity between 30 and 70%.

For *in vivo* triple-negative breast cancer modeling and CAR-T treatment, 3 x 10^6^ HCC70-GFP cells mixed with 3 x 10^6^ TNBC-derived CAFs in 50 μL of ice-cold PBS: Matrigel (1:1) were injected into the mammary fat pad of 8-weeks-old, female NSG mice. Mice were randomly enrolled into the study once tumor volume reached ~50 mm3. For CAR T-cell treatment, tumor bearing mice received a single-dose treatment of mock-transduced T-cells (5 x 10^6^ cells/mouse), FAP UCAR T-cells (5 x 10^6^ cells/mouse) or Mesothelin UCAR T-cells (3 x 10^6^ cells/mouse) in 100□μl. of PBS via intravenous injection. Where indicated, mice were treated with 10 mg/kg anti-hPD-1 (BioXCell, Catalog No. SIM0003) via intra-peritoneal injection, once every other day over a period of 6 days. The mice were monitored for health, weighed at least once weekly, and followed to measure survival. Disease progression was monitored on a weekly basis by measuring tumor dimensions using digital calipers and tumor volume was calculated using the formula [length x (width)2/2]. Humane endpoint criteria for tumor models were (i), weight loss greater than or equal to 20% from baseline, (ii), abnormal gait, paralysis, or inability to ambulate properly, (iii), respiratory distress/labored breathing, (iv), lethargy or persistent recumbency, and (v), loss of righting reflex or other abnormal neurological behaviors. The method for euthanasia was CO2 asphyxiation followed by cervical dislocation to assure death.

### Histology and IHC

Mouse tumors were fixed in 10% buffered formalin (Thermo Fisher Scientific) overnight and moved to 70% ethanol thereafter. Paraffin embedding, tissue sectioning, H&E staining, Immunohistochemistry (IHC) and Trichrome staining was performed at Histowiz Inc., NY. For IHC antibodies used were: α-CD8, #LSB3914, LSBio and α-FAP, #ab227703, Abcam.

## Results

### Engineering non-alloreactive, immune evasive CAR modified T-cells targeting desmoplastic solid tumors

Universal ‘off-the shelf’ CAR T-cells circumvent several disadvantages of the autologous approach, including superior manufacturing efficiency with low cost and production time as well as the ability to use healthy donor T-cells with high potency and fitness^34^. Thus, using a combination of previously described TAL Effector Nucleases (TALEN) and recombinant lentiviral CAR particles, we engineered non-alloreactive CAR T-cells targeting the tumor cell antigen Mesothelin (derived from mesothelin antibody clone MN) and stromal CAF cell antigen FAP (**Figure 1A**)^35^. Two FAP-CAR constructs derived from previously described scFvs were tested in this study-hereby referred to as the anti-murine mF3^36^ which cross-reacts with human FAP protein and the humanized hF1. Human primary T-cells from healthy donors were transduced and TALEN transfected as outlined in **Figure 1B.** The engineering strategy includes two TALEN-mediated knockouts to generate non-alloreactive, immune evasive CAR-modified T-cells: TRAC, to prevent Graft-versus-Host disease, and B2M to downregulate surface HLA-ABC expression which confers resistance to Host-versus-Graft rejection, thereby increasing their persistence in an allogeneic context (**Figure 1C**). Robust gene-edited CAR T-cell engineering was confirmed by flow cytometry with significant expression of all three CAR constructs relative to mock transduced control (~40-60%), and efficient TALEN gene editing, with upwards of 80% TCRα/β and HLA-ABC double knock-out cells (**Figure 1D**). Henceforth for the sake of clarity, TCRα/β and HLA-ABC double knock-out T-cells will be referred to as Universal T-cells. Mock transduced Universal T-cells, FAP CAR T-cells and Mesothelin CAR T-cells will be referred to as UT, FAP UCAR and Meso UCAR T-cells, respectively.

**FIGURE 1.**
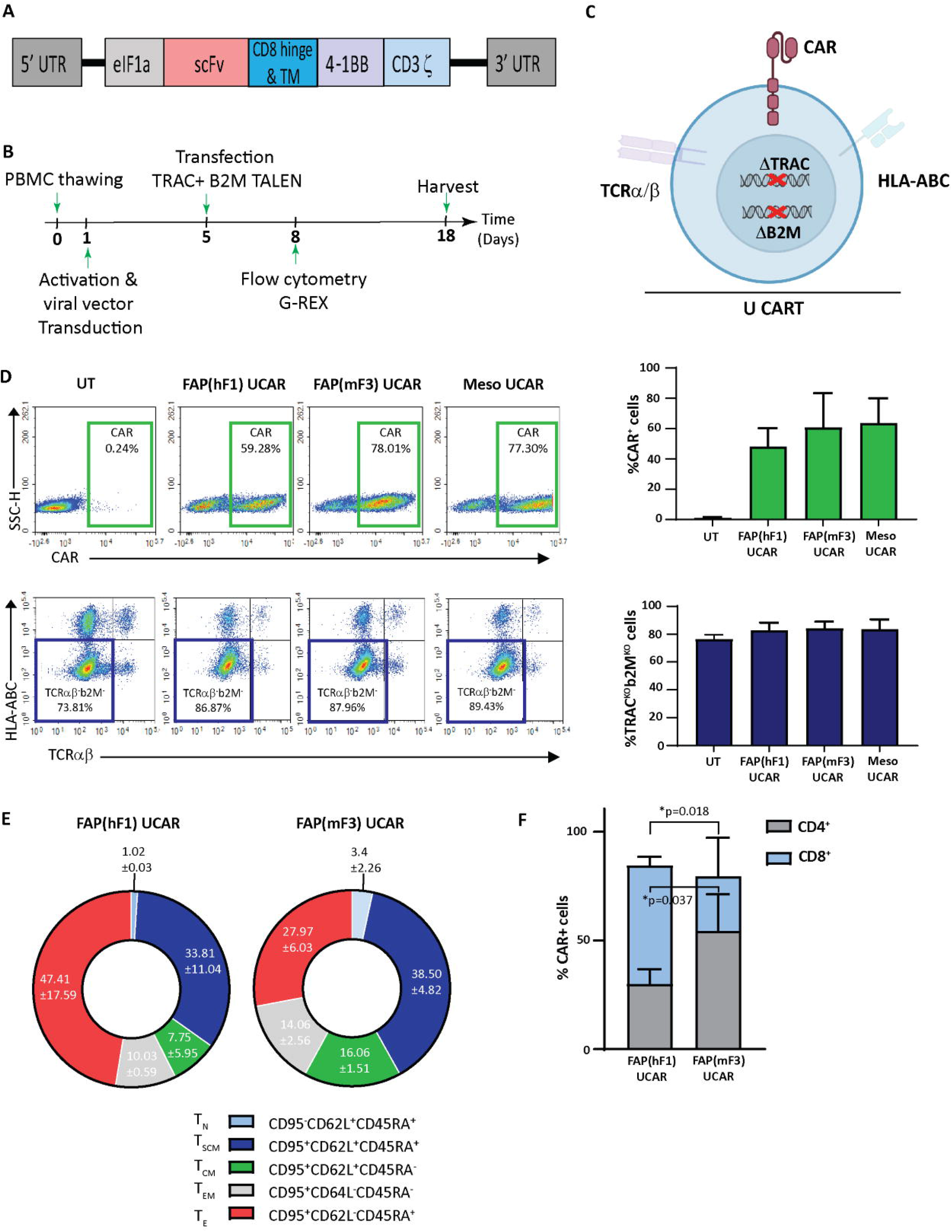
TALEN-mediated multiplex editing generates universal immune evasive CAR T-cells against solid tumor targets. **(A)** Retroviral CAR construct for T cell transduction comprised of the anti-tumor targeting single-chain variable fragment (scFv), human CD8α hinge and transmembrane domain, a human 4-1BB costimulatory domain, and a human CD3ζ activation domain. **(B)** Experimental strategy for TALEN-mediated gene editing, lentiviral transduction, expansion, and analysis of engineered human universal CAR T-cells. **(C)** Pictorial representation of immune evasive, universal CAR T-cells engineered with lentiviral CAR expression and TALEN-mediated multiplex editing of TRAC and B2M gene loci, resulting in downregulation of surface TCRα/β and HLA-ABC. **(D)** Top panel, flowcytometry plots showing frequency of CAR expression among viable engineered T cells (left), and associated quantitation (right). Bars show the means ± SD, n=3. Bottom panel, flow-cytometry plots showing frequency of TCRα/β (-)/HLA-ABC (-) viable engineered T cells (left) and associated quantitation (right). Bars show means ± SD, n=3. **(E)** Frequency of FAP UCAR T-cell subpopulations displaying CD95^-^CD62L^+^CD45RA^+^ (T_N_ naive-like), CD95^+^CD62L^+^CD45RA^+^ (T_SCM_ memory stem cell), CD62L^+^CD45RA^-^ (T_CM_ central memory), CD62L^-^CD45RA^-^ (T_EM_ effector memory) and CD62L^-^CD45RA^+^ (T_E_ terminal effector) labeling at end of engineered T cell expansion (n=2). **(F)** Frequency of FAP UCAR T-cell CD4^+^ and CD8^+^ subpopulations (n=3). Bars show the means ± SD; P-values determined by Student *t* test (two-tailed, unpaired).

The two different FAP UCAR T-cell populations were further characterized by flow cytometry to determine variance in their cell differentiation states and CD8^+^/CD4^+^ states, which could have potential functional consequences. Cell differentiation states of FAP(hF1) UCAR and FAP(mF3) UCAR T-cells were comparable, with the effector T-cell (T_E_) frequency higher in the former than the latter, though not significantly (**Figure 1E**, **Figure S1A**). Remarkably, FAP(hF1) UCAR T-cells had significantly higher frequency of CAR^+^ CD8^+^ cells relative to FAP(mF3) UCAR T-cells, which could be indicative of a higher response rate *in vivo*^38^ (**Figure 1F**, **Figure S1B**).

### Meso CAR T-cell anti-tumor cytotoxicity is enhanced in combination with CAF-targeting FAP CAR T-cells *ex vivo*

To comparatively assess the cytotoxic activity of FAP UCAR T-cells, FAP-expressing CAFs isolated from patient TNBC tumor (**Figure S2A**) were used as targets in an *in vitro* cytotoxic assay, outlined in **Figure 2A**. Both FAP(hF1) UCAR and FAP(mF3) UCAR T-cells killed TNBC-derived CAFs with equal efficiency (**Figure 2B**). Interestingly though, when we assessed IFNγ release by these two CAR T populations when co-incubated with FAP^+^ CAFs, FAP(hF1) UCAR T-cells secreted significantly higher amounts of IFNγ, relative to FAP(mF3) UCAR T-cells (**Figure 2C**).

**FIGURE 2.**
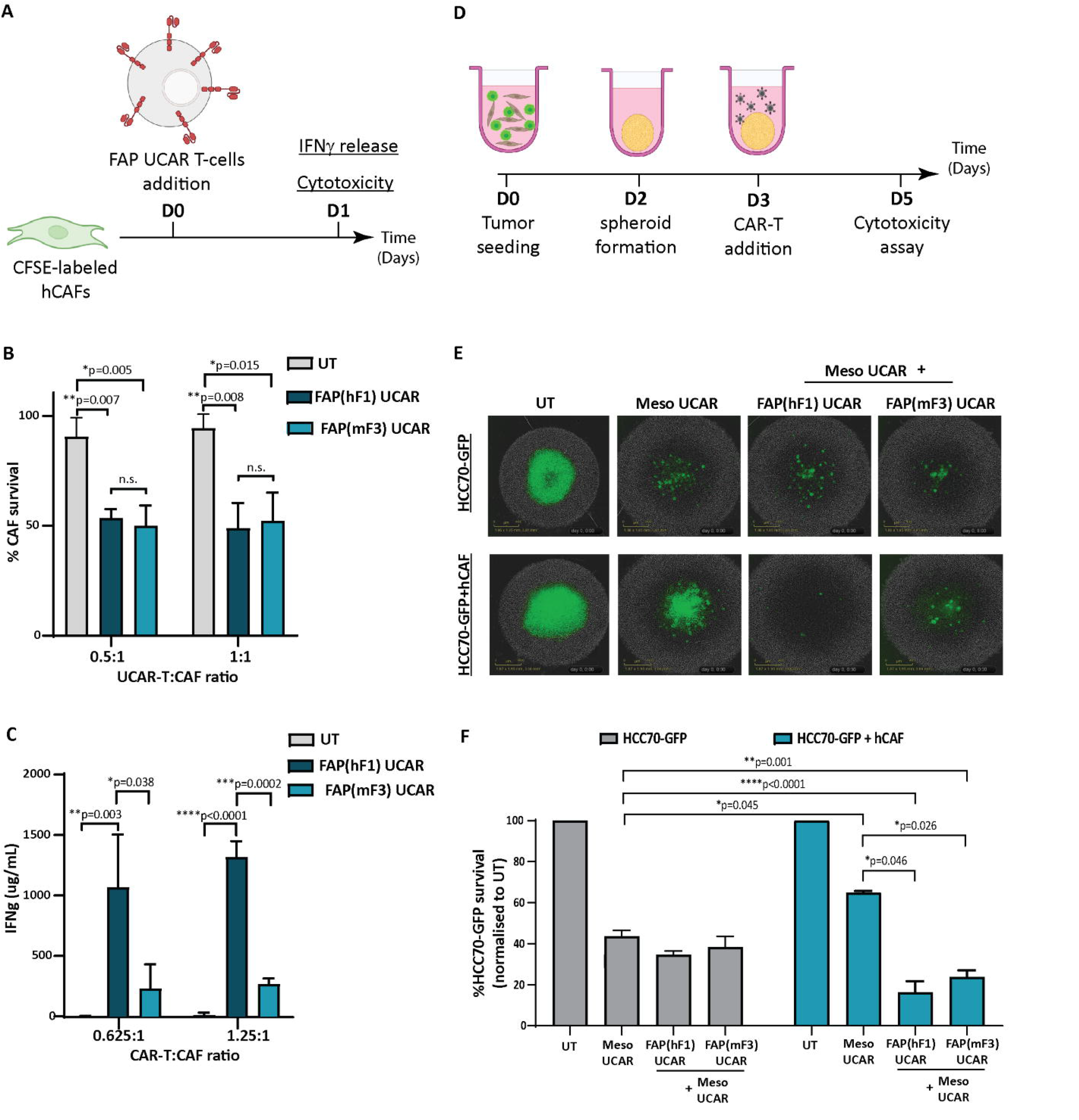
FAP UCAR T-cells efficiently kill FAP^+^ CAFs and enhance Meso UCAR T-cell cytotoxicity against tumor cells and CAF spheroid co-cultures *in vitro*. **(A)** Experimental strategy for assessing FAP UCAR T-cell IFNγ-release and cytotoxicity against TNBC patient-derived CAFs. **(B)** Bar graph representing percentage of CAF survival post 24 hour co-incubation with UT, FAP(hF1) UCAR or FAP(mF3) UCAR T-cells at two different Effector:Target ratios. Results representative of two independent experiments with two donors each, n=3 technical replicates per donor per experiment. Bars show the means ± SD; P-values determined by Student *t* test (two-tailed, unpaired). **(C)** Bar graph representing IFNγ levels in supernatant post 24-hour co-incubation of CAF with UT, FAP(hF1) UCAR or FAP(mF3) UCAR T-cells at two different Effector: Target ratios. Results representative of two independent experiments with two donors each, n=4 technical replicates per donor per experiment. Bars show the means ± SD; P-values determined by Student *t* test (two-tailed, unpaired). **(D)** Schematic of FAP UCAR and Meso UCAR T-cell cytotoxicity assay against 3-D spheroids of TNBC cell line HCC70-GFP alone or co-cultured with TNBC-derived CAFs. **(E)** Representative fluorescent images of HCC70-GFP and HCC70-GFP co-incubated with TNBC-derived CAF spheroids on day 5 of cytotoxicity assay as outlined in **(D)**. **(F)**Bar graph representing percentage HCC70-GFP tumor cell survival post cytotoxicity assay outlined in **(D)**. Data representative of two independent experiments with two donors each, n=4 technical replicates per donor per experiment. Bars show the means ± SD; P-values determined by Student *t* test (two-tailed, unpaired).

As described above, CAFs are critical drivers of a desmoplastic tumor microenvironment which in turn downregulates T-cell infiltration in these tumors. Thus, following confirmation of anti-CAF cytotoxicity of both FAP CAR T-cells, we next assessed their ability to make desmoplastic ‘cold’ solid tumors more susceptible to tumor antigen-directed CAR T-cells. For this purpose, we generated physiologically relevant *ex vivo* heterotypic TNBC spheroids composed of mesothelin-expressing TNBC cell line HCC70 expressing a GFP reporter (HCC70-GFP) (**Figure S2B**) and TNBC-derived CAFs at 1:2 ratio. Homotypic TNBC spheroids composed of HCC70-GFP cells alone were used as controls in a CAR T cytotoxicity assay outlined in **Figure 2D**. As indicated in **Figure 2E** and **Figure 2F**, while HCC70-GFP alone spheroids were vulnerable to Meso UCAR T-cell cytotoxicity, CAF addition to these tumor spheroids significantly decreased Meso UCAR T-cell cytotoxicity. Remarkably, combination treatment of Meso UCAR T-cells with both FAP(hF1) and FAP(mF3) UCAR T-cells not only restored but further enhanced anti-tumor activity against the HCC70+CAF spheroids significantly, relative to Meso UCAR T-cell activity against HCC70 spheroid alone (**Figure 2F**). Altogether our results indicate that while the presence of CAFs in *ex vivo* TNBC spheroids makes them resistant to Meso UCAR T-cell cytotoxicity, this resistance can be circumvented by concurrent targeting of CAFs by FAP UCAR T-cells.

### CAF ablation by FAP UCAR T-cells reduces desmoplasia and promotes T-cell infiltration and anti-tumor activity *in vivo*

In the interest of potential clinical applications, it was essential to validate the activity of human allogeneic CAR T-cells in a physiologically relevant *in vivo* solid tumor model, with distinct tumor and stromal compartments. Pursuant to this, we orthotopically implanted TNBC cell line HCC70-GFP in combination with patient TNBC tumor-derived CAFs in NSG mice mammary fat pad (**Figure S3A**). As previously reported^39^, co-implantation of HCC70 cells with CAFs in different ratios promoted the rate of tumor progression, with the most significant cooperative growth observed for HCC70 and CAFs co-implanted at 1:1 ratio (**Figure S3B**). Successful implantation of injected CAFs in these tumors was confirmed by positive staining of spindle-like cells with human-specific FAP antibody (**Figure S3C**). Going forward, we employed this orthotopic model of TNBC for further investigations.

We commenced by dissecting the activity of FAP(hF1) UCAR, FAP(mF3) UCAR and Meso UCAR T-cells against orthotopic TNBC tumors, as outlined in **Figure 3A**. FAP(hF1) UCAR T-cells alone significantly reduced tumor growth (**Figure 3B**) relative to UT-cell controls. Immune profiling of harvested tumors revealed robust tumor infiltration and accumulation of FAP(hF1) UCAR T-cells, as measured by quantity of detected human CD45^+^ cells (**Figure 3C** and **Figure 3D**). The human CD45^+^ population was further analysed for CD4^+^/CD8^+^ distribution, with significantly higher CD8^+^ sub-populations in the FAP(hF1) UCAR and FAP(mF3) UCAR groups. (**Figure 3E**). Significantly higher CD8^+^ cells were also detected by immunohistochemistry in tumors treated with FAP(hF1) UCAR T-cells, relative to all other groups (**Figures 3F**, **3G**). Furthermore, while FAP(hF1) UCAR T-cells were detected in the spleen of mice treated with these cells (**Figure S3D)**, neither FAP(mF3) UCAR nor Meso UCAR T-cells were detectable in respectively treated mouse splenocytes. This suggested that while FAP(hF1) UCAR T-cells displayed activation-induced systemic accumulation, FAP(mF3) and Meso UCAR T-cells failed to do so.

**FIGURE 3.**
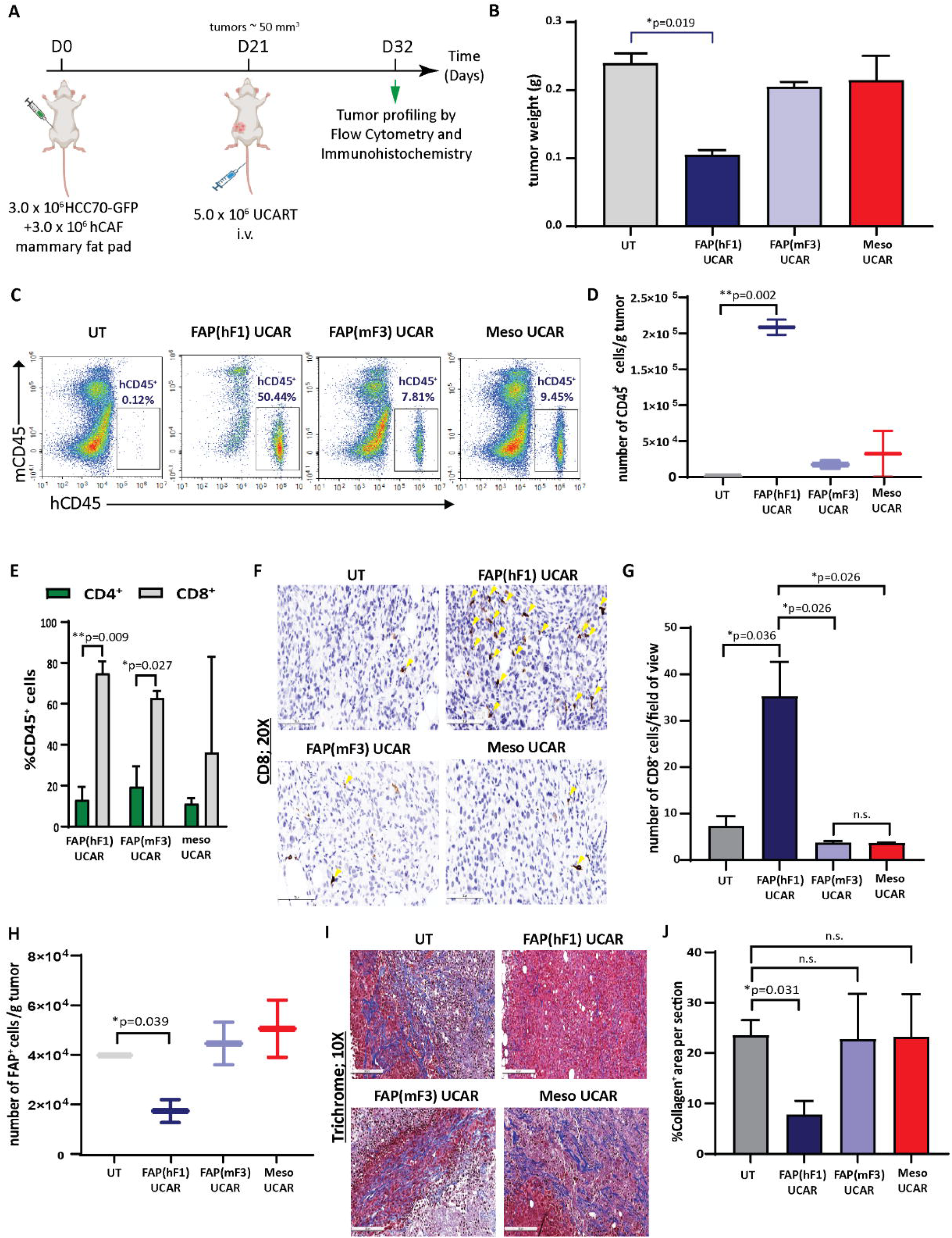
FAP(hF1) UCAR T-cells successfully deplete CAFs and accumulate in TNBC tumor stroma *in vivo* **(A)** Schematic of UCAR T-cell treatment and analysis of orthotopic TNBC tumors implanted in NSG mice. **(B)** Bar graph representing weights of tumor harvested from mice treated with control UT, FAP(hF1) UCAR, FAP(mF3) UCAR or Meso UCAR T-cells as per schematic in **(A).** Bars show the means ± SEM; P-values determined by Student *t* test (two-tailed, unpaired), n=2. **(C)** Flow cytometry plot of human CD45^+^ UCAR T-cell population in orthotopic TNBC tumors harvested from mice treated as indicated. **(D)** Box and whiskers plot representing quantitation of total number of CD45^+^ cells per gram tumor, calculated using **(C).** P-values determined by Student *t* test (two-tailed, unpaired), n=2. **(E)** Stacked bars plot representing CD4^+^ and CD8^+^ population, as percentage of total human CD45^+^ cells, in orthotopic TNBC tumors from mice treated with indicated UCAR T-cells. Bars show the means ± SEM, n=2. **(F)** Immunohistochemistry for detection of human CD8^+^ cells in orthotopic TNBC tumors from mice treated with indicated UCAR T-cells. Yellow arrows indicate human CD8 stained cells. **(G)** Bar graph representing quantitation of **(F)**, each section divided into 4 fields of view. Bars show the means ± SEM; P-values determined by Student *t* test (two-tailed, unpaired), n=2. **(H)** Box and whiskers plot representing quantitation of total number of human FAP^+^ cells per gram tumor, as determined by flow cytometry analysis of orthotopic TNBC tumors from mice treated with indicated UCAR T-cells. P-values determined by Student *t* test (two-tailed, unpaired), n=2. **(I)** Trichrome staining of orthotopic TNBC tumor sections from mice treated with indicated UCAR T-cells. Collagen network indicated by blue staining. **(J)** Bar graph representing quantitation of blue stained Collagen area per section **(F)**, measured using ImageJ. Bars show the means ± SEM; P-values determined by Student *t* test (two-tailed, unpaired), n=2.

Functionally, high intra-tumoral FAP(hF1) UCAR T-cell levels positively correlated with significant depletion of FAP^+^ cells in these tumors (**Figure 3H, Figure S3E)**). This was accompanied by reduced desmoplasia in FAP(hF1) UCAR T-cells treated tumor stroma, as measured by collagen depletion (**Figures 3I, 3J**). In sharp contrast, FAP(mF3) or Meso UCAR T-cell-treated tumors populated with FAP^+^ CAFs (**Figure 3H**) were highly desmoplastic and sparsely infiltrated with CD45^+^ (**Figure 3C**) and CD8^+^ T-cells (**Figures 3F, G**). Taken together our results indicate that FAP(hF1) UCAR T-cell treatment of tumor-bearing mice results in significant FAP^+^ cells depletion and reduction in stromal physical barrier in the tumor microenvironment, confirming their onsite anti-CAF activity. This is accompanied by an increase in intra-tumor CD8^+^ FAP(hF1) UCAR T-cell levels and a substantial reduction in tumor growth, consistent with loss of tumor-supportive CAFs.

### FAP UCAR T-cell treatment sensitizes ‘cold’ desmoplastic tumors to subsequent Meso UCAR T-cell infiltration and anti-tumor cytotoxicity

Since FAP(hF1) UCAR T-cell-mediated CAF depletion was accompanied by reduced collagen deposition and increased tumor-infiltrating CAR T-cells, we postulated that this re-programming of the tumor microenvironment could make it more amenable to subsequent infiltration and cytotoxicity of otherwise inactive tumor-targeting CAR T-cells. To test this hypothesis, we designed an *in vivo* study as outlined in **Figure 4A,** wherein orthotopic TNBC-bearing NSG mice were treated with Meso UCAR T-cells either alone or subsequent to FAP(hF1) UCAR T-cell treatment. While Meso UCAR or FAP(hF1) UCAR T-cell treatment alone did not affect tumor growth, relative to UT-cell control, FAP(hF1) UCAR T-cells pre-treatment with subsequent Meso UCAR T-cell administration significantly decreased tumor size, relative to all other treatment groups (**Figure 4B, Figure S4A**). Cellular profiling of harvested tumors revealed highest human CD45^+^ levels in the FAP(hF1) UCAR T-cell plus Meso UCAR T-cell combination treatment group (**Figure 4C**). More importantly, significantly high levels of Meso UCAR T-cells were detected in FAP(hF1) UCAR T-cell pre-treatment group (**Figure 4D, Figure S4B**), which corresponded with maximal FAP^+^ CAF depletion (**Figure 4E, Figure S4C**). Taken together, these results validate the advantage of FAP UCAR T-cell pre-treatment of CAF-rich cold tumors to increase infiltration of subsequently administered tumor-antigen CAR T-cells.

**FIGURE 4.**
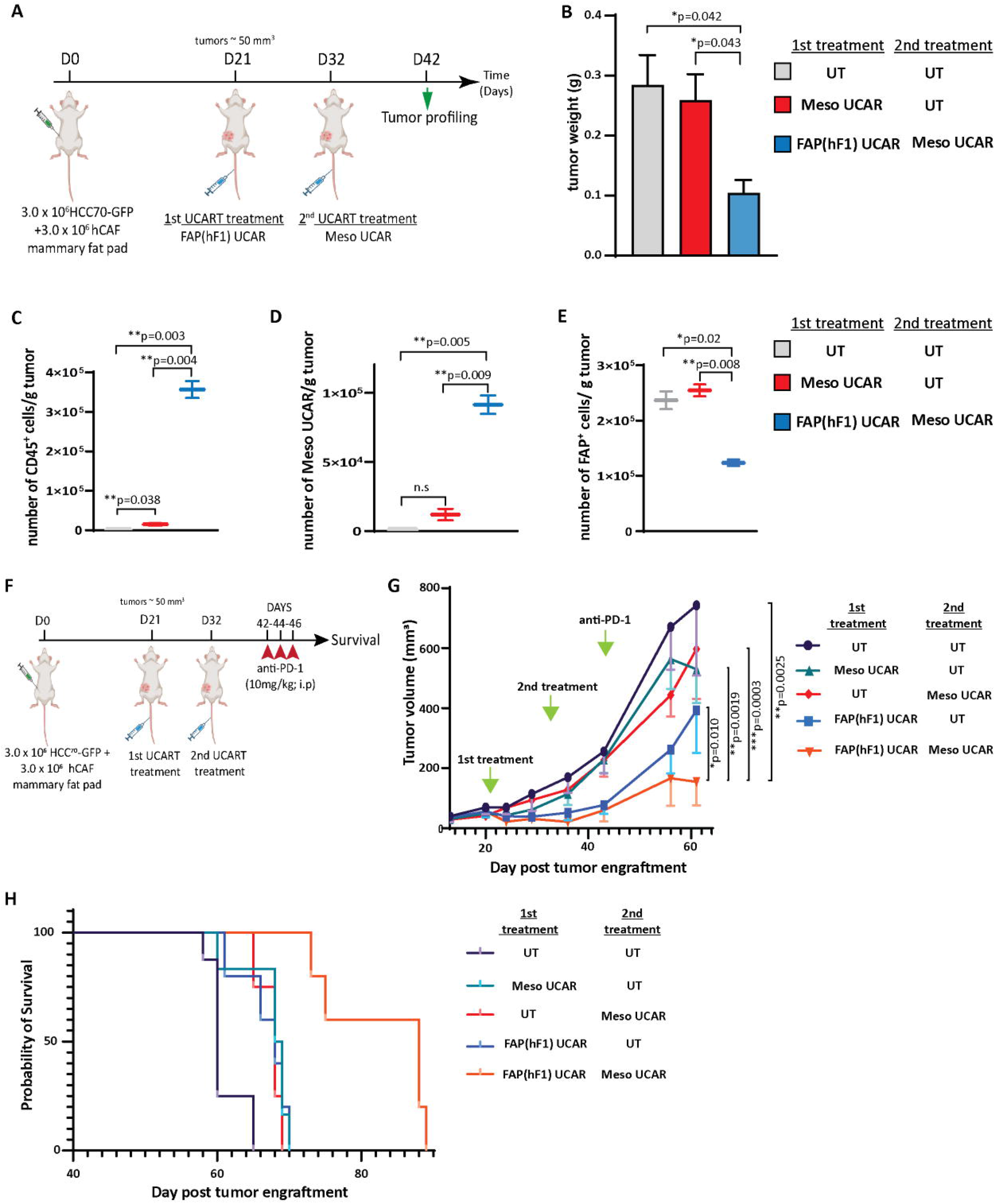
FAP(hF1) UCAR T-cell pre-treatment sensitizes CAF^+^ cold tumors to Meso UCAR T-cell plus anti-PD-1 combination therapy **(A)** Schematic of sequential UCAR T-cell treatment and analysis of orthotopic TNBC tumors implanted in NSG mice. **(B)** Bar graph representing weights of tumor harvested from mice treated with control UT, FAP(hF1) UCAR, or FAP(hF1) UCAR and Meso UCAR T-cells combination as per schematic in **(A).** Bars show the means ± SEM; P-values determined by Student *t* test (two-tailed, unpaired), n=2. **(C)** Box and whiskers plot representing quantitation of total number of CD45^+^ cells per gram orthotopic TNBC tumors from mice treated with indicated UCAR T-cells, as determined by flow cytometry. P-values determined by Student *t* test (two-tailed, unpaired), n=2. **(D)** Box and whiskers plot representing quantitation of total number of Meso UCAR T-cells per gram orthotopic TNBC tumors from mice treated with indicated UCAR T-cells, as determined by flow cytometry. P-values determined by Student *t* test (two-tailed, unpaired), n=2. **(E)** Box and whiskers plot representing quantitation of total number of FAP^+^ cells per gram orthotopic TNBC tumors from mice treated with indicated UCAR T-cells, as determined by flow cytometry. P-values determined by Student *t* test (two-tailed, unpaired), n=2. **(F)** Schematic of sequential UCAR T-cell and anti-PD-1checkpoint inhibitor treatment and subsequent analysis of orthotopic TNBC tumor-implanted NSG mice. **(G)** Graph representing growth kinetics of orthotopic TNBC tumors in mice treated as indicated over time. P-values determined by Student *t* test (two-tailed, unpaired), n=5-8 mice per cohort. **(H)** Kaplan-Meier curve for survival analysis of orthotopic TNBC tumor-implanted NSG mice treated as indicated (n=5-8 per cohort). P-values of Log-Rank test were as follows: UT vs Meso UCAR +UT **p=0.0026; UT vs UT+Meso UCAR **p=0.0060; UT vs FAP(hF1) UCAR + UT **p=0.0050; UT vs FAP(hF1) UCAR + Meso UCAR ***p=0.0008.

Furthermore, we designed a long-term combination treatment study to determine tumor growth and survival kinetics of the strategy **(Figure 4F**). Consistent with promising ongoing clinical trials using mesothelin CAR T-cells in combination with PD-1 checkpoint blockade^40, 41^, we included an anti-PD-1 (nivolumab mimetic) co-treatment arm, to determine whether FAP UCAR T-cell addition to this combination can further enhance efficacy. FAP(hF1) UCAR T-cell pre-treatment with subsequent Meso UCAR T-cell administration exerted efficient tumor growth inhibition and conferred highest survival benefit to tumor-bearing mice compared to the other experimental groups (**Figures 4G-H**). Indeed, tumor growth inhibition and survival benefit obtained with Meso UCAR T-cell or FAP(hF1) UCAR T-cell treatments alone were significantly lower than with the two combined. It is to be noted however that, although not significant, FAP(hF1) UCAR T-cells alone did decrease tumor growth, compared to UT-cell group (p=0.051, **Figure 4G**). In contrast, while active *in vitro*, FAP(mF3) UCAR T cells also failed to sensitize these tumors to Meso UCAR T-cell cytotoxicity *in vivo* (**Figures S4D,S4E**), in agreement with their inability to deplete CAFs or increase CD8^+^ T-cell infiltration in the tumor stroma (**Figures 3H-J**).

## Discussion

In this study, we conceived and validated a novel immunotherapeutic combination strategy aimed at sensitizing CAF positive ‘cold’ tumors to tumor antigen-targeting CAR T-cell infiltration and cytotoxicity. Using TALEN-mediated gene editing, we engineered universal hypoimmunogenic CAR T-cells targeting components of a TNBC solid tumor: FAP UCAR T-cells against stromal CAFs and Meso UCAR T-cells against tumor cells. Our *in vitro* and *in vivo* results show that FAP UCAR T-cells enable the reprogramming of the cold, stroma-rich TNBC TME, making the tumor susceptible to subsequent Meso UCAR T infiltration and cytotoxicity and improving the overall antitumor activity of the treatment.

While there are several reports evaluating FAP-targeting CAR T-cell strategies for CAF depletion in the solid TME^36, 37, 42–45^, what is evident from these studies is that (i) the ‘on-target off-tumor’ cytotoxicity of FAP CAR T-cells is probably FAP scFv specific and choosing the appropriate scFv may alleviate this safety concern, and (ii) while efficiently ablating TME CAFs, FAP CAR-T treatment alone displays limited anti-tumor activity. For instance, while Tran et al^44^ observed ‘on target off-tumor’ cytotoxicity in mouse bone marrow using FAP5 scFv, no such toxicities have been reported in other studies which use different anti-FAP scFvs^36, 37, 43, 45^. Thus, our two main objectives were (i) to select a FAP-scFv which in our selected CAR domain scaffold (**Fig. 1a**) and as an TALEN-gene edited CAR T-cell is safe and effective against human CAFs *in vivo* and (ii) leverage the ability of FAP CAR T-cells to turn ‘cold’ tumors ‘hot’ to develop combination immunotherapy strategies with higher anti-tumor activity.

To test our strategy, we optimized an orthotopic TNBC model in NSG mice using both patient tumor-derived CAFs and tumor cell line, to generate a physiologically relevant system that closely recapitulates one of the biggest hurdles to CAR-T therapeutic success, namely tumor homing and T cell infiltration. Furthermore, the model allowed us to assess the de facto allogeneic CAR T-cells that will translate from bench-to-bedside if found safe and effective. To our knowledge, this is the first report demonstrating pre-clinical activity of non-alloreactive human CAR T-cells against an orthotopic TNBC tumor with humanized stroma. Our validated FAP(hF1) UCAR T-cell anti-tumor activity concurs with the observations of Schuberth et al in a mesothelioma xenograft model with tumor cell FAP expression^37^ and was further determined to be safely tolerated in a phase I clinical trial of autologous FAP(hF1) CAR T-cells against malignant pleural mesothelioma (MPM)^40^. Thus, we believe that FAP(hF1) UCAR T cells hold great promise for clinical development against solid tumors.

Additionally, we demonstrate the effectiveness of pre-treating stromal solid tumors with FAP(hF1) UCAR T cells before administering tumor-antigen targeting CAR T-cells, in this instance Meso UCAR T-cells. While multiple clinical trials have demonstrated a favorable safety profile for Mesothelin CAR T-cells, either alone or in combination with PD-1 checkpoint inhibition, therapeutic efficacy against most solid tumors remains modest^40, 41, 46, 47^. In agreement, we observed that patient-derived CAF^+^ orthotopic TNBC tumors were largely unresponsive to Meso UCAR T cells and marked by poor CAR T-cell tumor infiltration, even in the presence of immune checkpoint inhibition (ICI). However, depletion of these CAFs by FAP(hF1) UCAR T-cells decreased desmoplasia and enabled subsequently administered Meso UCAR T-cells to infiltrate the tumor. This combination strategy markedly reduced primary tumor progression and significantly extended mice survival, thus validating the therapeutic advantage of FAP CAR T-cell treatment prior to tumor antigen targeting CAR T-cell administration for stromal solid tumors. The mice eventually succumbed to the disease, an outcome that could be attributable to tumor metastasis since secondary tumors were detected in these mice, predominantly in the lung. Optimizing CAR T-cell dosage and regimen as well as re-dosing can be potential methods of improving treatment efficacy.

Since FAP(hF1) UCAR T-cell treatment makes the TME more permissible to T cell infiltration, an additional application of these cells in a therapeutic strategy is to sensitize previously unresponsive ‘cold’ tumors to checkpoint inhibition. While immune checkpoint inhibitors have shown remarkable response in a variety of solid tumors, there remains a large patient population unresponsive to this therapy. Primary resistance to ICI in a sub-set of patients correlates with poor tumor-infiltrating lymphocytes (TILs), while majority ICI responders have high TIL levels^49^. We thus postulate FAP(hF1) UCAR T-cell treatment can potentially increase intra-tumoral T cell infiltration in patients with cold tumors, thus increasing their probability to respond positively to subsequent ICI therapy. Since TILs are composed of a spectrum of antigen specificity, this has an added advantage of addressing the complication of solid tumor heterogeneity, leading to more efficient tumor clearance and preventing CAR antigen escape.

A potential hindrance to this approach is the requirement for lymphodepletion prior to allogeneic CAR T-cell administration, which can significantly hamper patient T-cell tumor infiltration. To circumvent this complication, our universal FAP UCAR T-cells are TALEN-edited to knockout beta-2 microglobulin (B2M), a strategy that allows the cells to evade host versus graft (HvG) rejection and extend their persistence in an allogeneic setting^35^. Our strategy thus doesn’t require long-lasting lymphodepleting agents such as Alemtuzumab, which can profoundly deplete endogenous immune effectors. Furthermore, unlike tumor cells, CAFs are primary cells with limited proliferative capacity. Thus, FAP(hF1) UCAR T-cell treatment should create a long-lasting CAF-depleted tumor milieu, allowing a window for reconstitution of the endogenous immune system and subsequent TIL activity in this CAF-depleted state. Furthermore, due to their the immune-evasive property, FAP(hF1) UCAR T cells can be readily combined with autologous tumor-targeting CAR T-cell therapies as well. Overall, we propose a combination treatment paradigm for solid tumors wherein pre-treatment of CAF positive tumors with FAP(hF1) UCAR T cells reprograms the TME from an immune desert to a T cell rich immune reactive milieu, thereby sensitizing them to other T cell-mediated immunotherapeutic modules.

## Author Contributions

**Shipra Das**: conceptualization, experimental design and execution, data analysis, figure preparation, manuscript drafting, reviewing, and editing. **J. Valton**: conceptualization, experimental design, manuscript reviewing and editing. **P. Duchateau**: conceptualization, manuscript reviewing and editing. **L. Poirot**: conceptualization, experimental design, data analysis, manuscript reviewing and editing.

## Acknowledgements

We would like to thank the clinical, translational and process and analytical development teams at Cellectis for their support through this work. We would also like to thank Cellectis publications team for their valuable advice.

