## Supplemental Material for "Stromal depletion by TALEN-edited universal hypoimmunogenic FAP-CAR T cells enables infiltration and anti-tumor cytotoxicity of tumor antigen-targeted CAR-T immunotherapy"

### Supplementary Figures

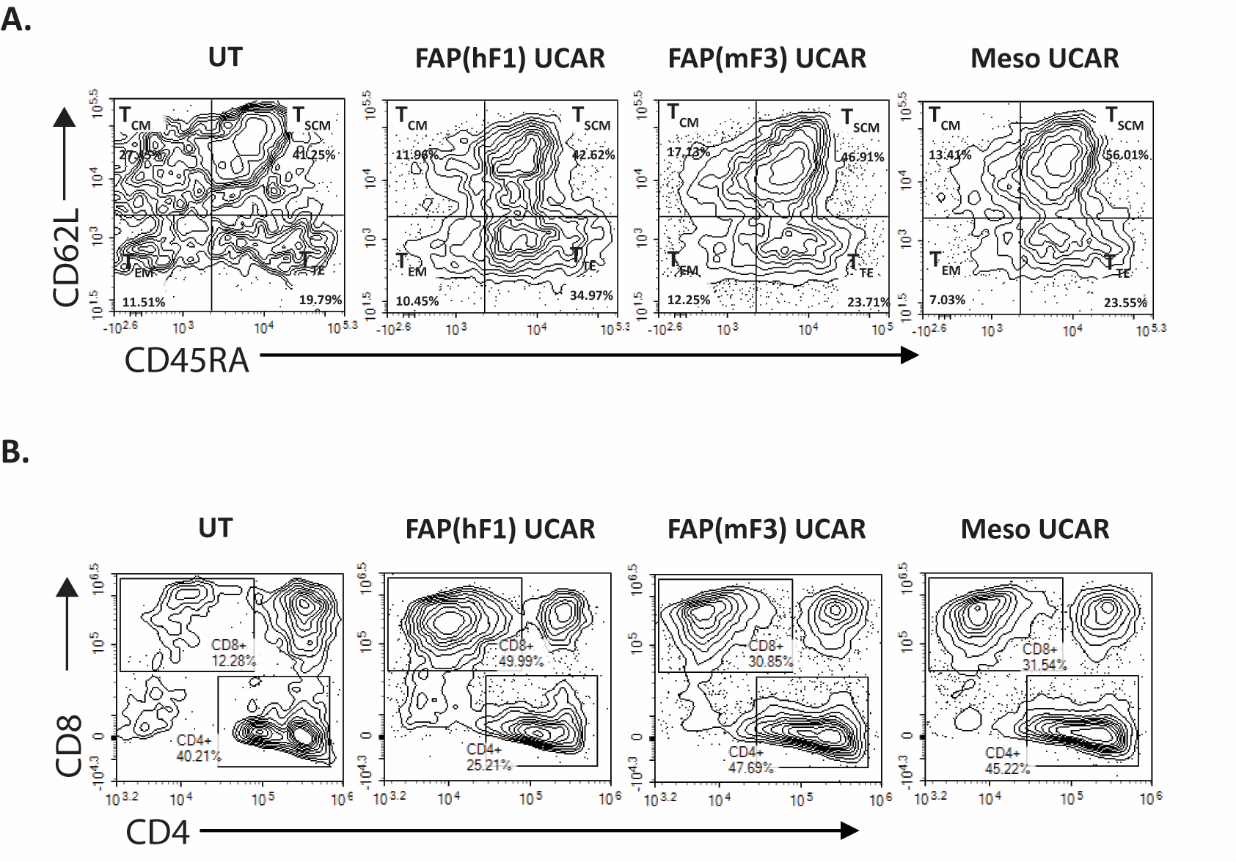

**Figure S1. (A)** Flow cytometry plots of T cell differentiated states of UCAR T-cells as indicated, pre-gated on viable, CD95^+^, CAR^+^ singlet cells or on viable, singlet cells for UT control. **(B)** Flow cytometry plots of CD4^+^ and CD8^+^ T-cell sub-populations of UCAR T-cells as indicated, pre-gated on viable, CAR^+^ singlet cells or on viable, singlet cells for UT control.

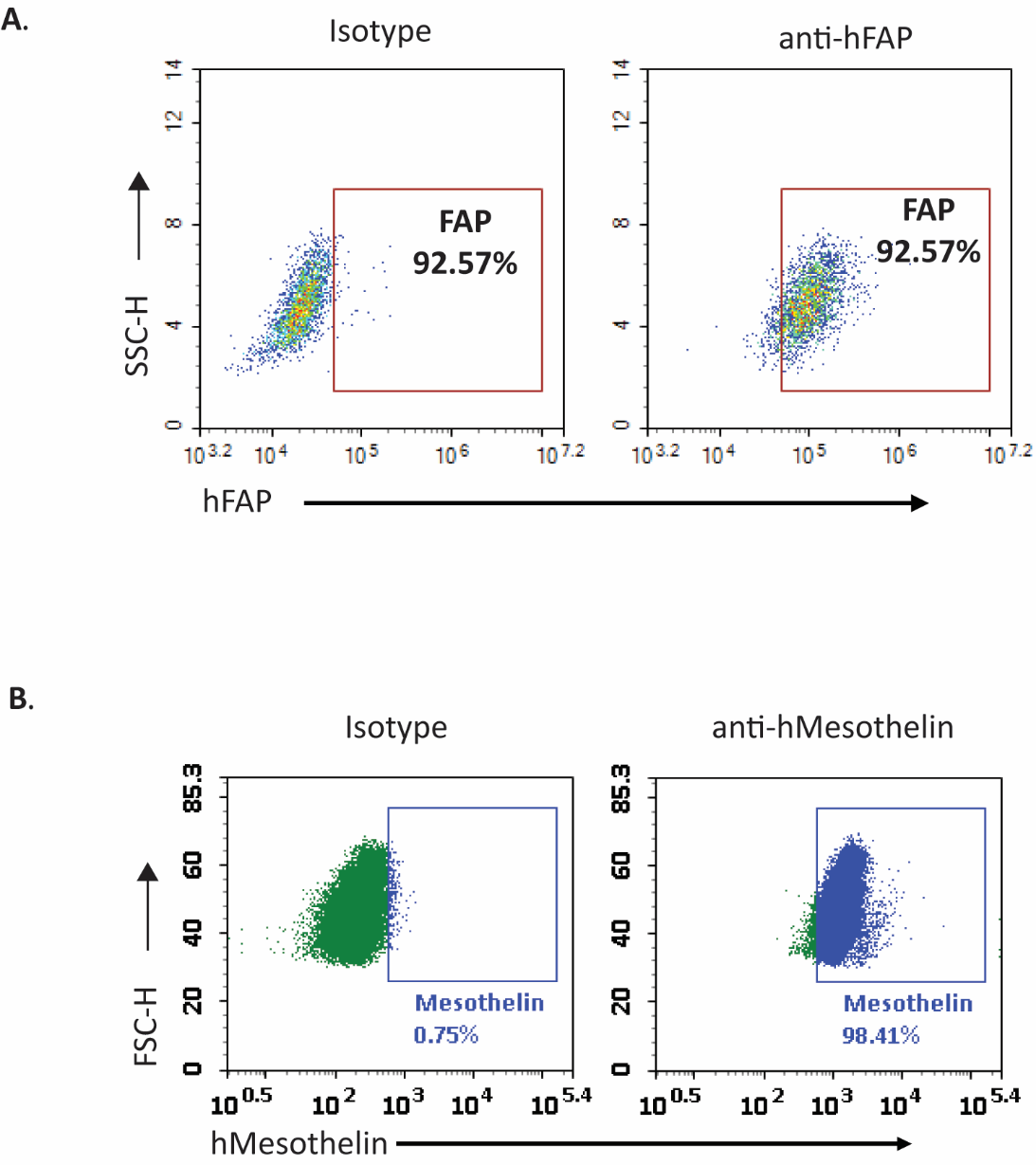

**Figure S2. (A)** Flow cytometry plots depicting staining of patient TNBC-derived CAF cells with isotype control or anti-human FAP protein antibody. **(B)**  Flow cytometry plots depicting staining of TNBC cell line HCC70-GFP with isotype control or anti- human Mesothelin protein antibody.

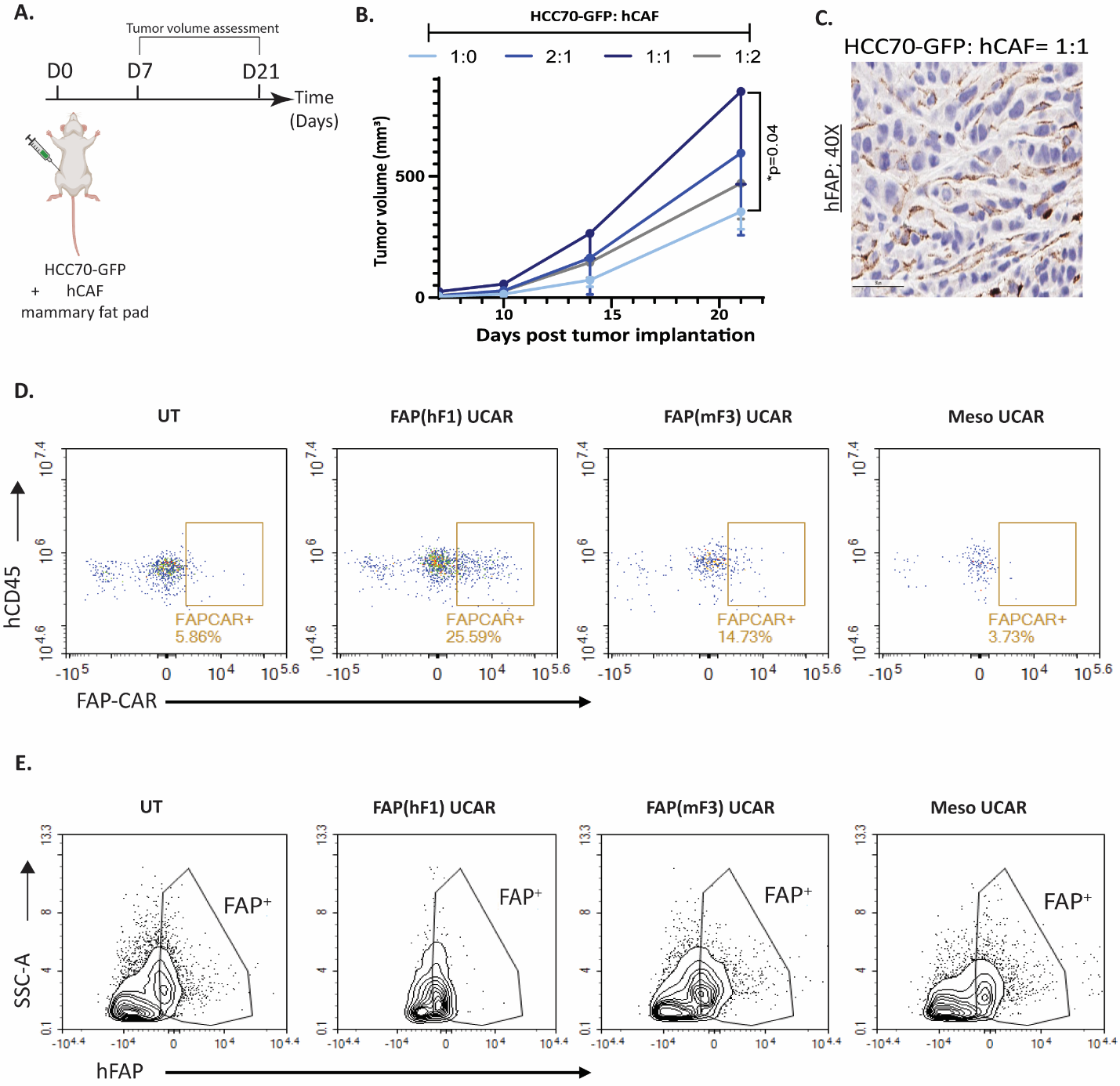

**Figure S3. (A)** Schematic depicting assessment of tumor growth kinetics upon co-implantation of 7 x 10^6^ HCC70-GFP cells with CAFs at different ratios in mammary fat pad of NSG mice. **(B)** Graph depicts growth kinetics of orthotopic mammary tumors implanted as outlined in **(A)**. **(C)** Immunohistochemical staining of human FAP protein in orthotopic tumor section derived from co-implantation of 7 x 10^6^ HCC70-GFP and 7 x 10^6^ TNBC-derived CAFs in mammary fat pad of NSG mice. **(D)** Flow cytometry plots depicting FAPCAR^+^ cells pre-gated on viable, mouse CD45^-^, human CD45^+^ singlet cells in spleen of orthotopic tumor bearing mice, treated as indicated. **(E)** Flow cytometry plots depicting human FAP^+^ cells pre-gated on viable, mouse CD45^-^, human CD45^-^ singlet cells in orthotopic tumors of NSG mice, treated as indicated.

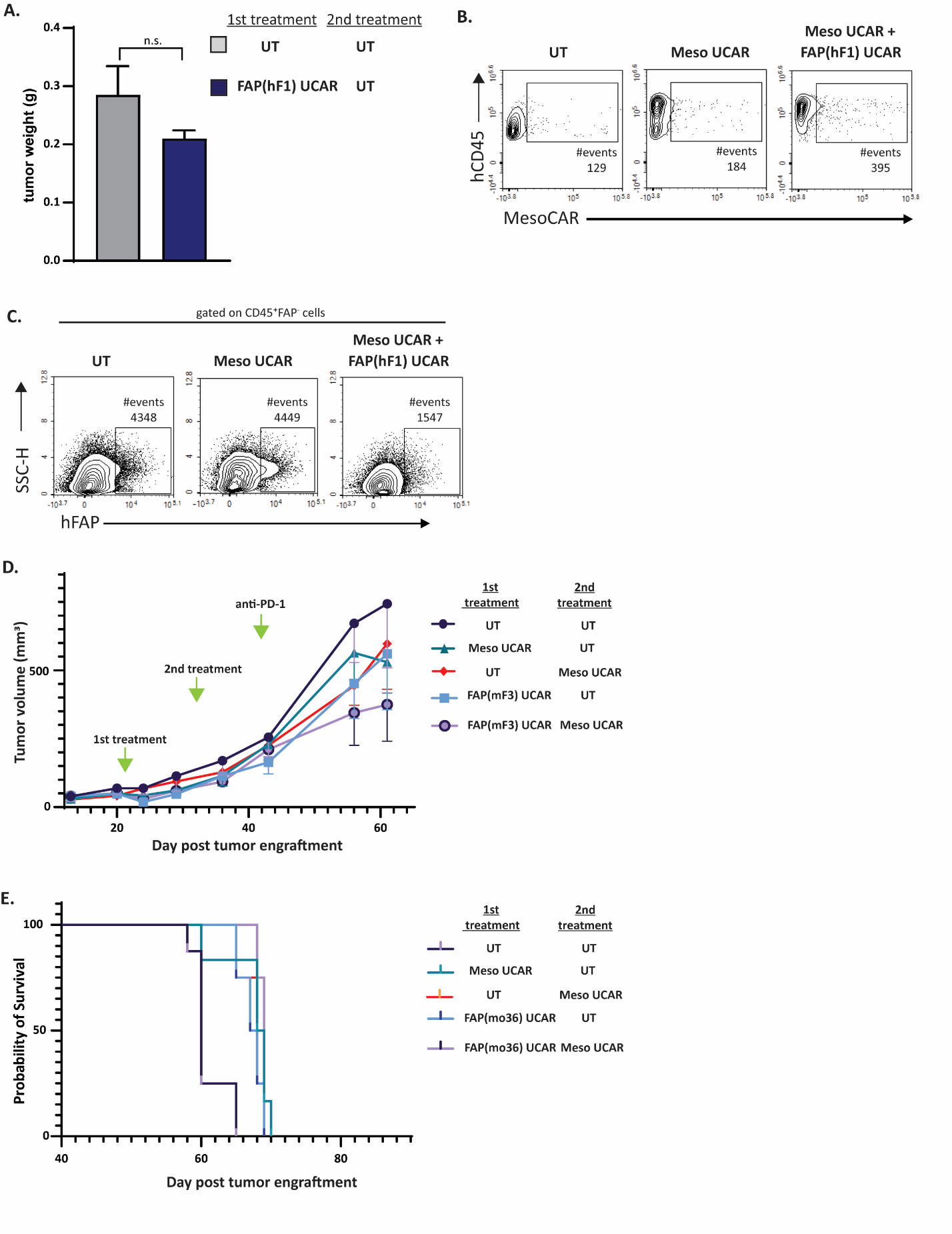

**Figure S4. (A)** Bar graph depicting tumor weight harvested from orthotopically injected NSG mice and treated with UCAR T-cells as indicated, 21 days post treatment initiation. n.s.-not significant. **(B)**  Flow cytometry plots depicting Mesothelin CAR^+^ cells pre-gated on viable, mouse CD45^-^, human CD45^+^ singlet cells in orthotopic tumors of NSG mice, treated as indicated. **(C)** Flow cytometry plots depicting human FAP^+^ cells pre-gated on viable, mouse CD45^-^, human CD45^-^ singlet cells in orthotopic tumors of NSG mice, treated as indicated. FAP^+^ gate was set using CD45^+^ cells which are negative for FAP. **(D)** Graph representing growth kinetics of orthotopic TNBC tumors in mice treated as indicated over time, n=5-8 mice per cohort. **(E)** Kaplan–Meier curve for survival analysis of orthotopic TNBC tumor-implanted NSG mice treated as indicated (n=5-8 per cohort).
